# Identification of potential key genes for SARS-CoV-2 infected human bronchial organoids based on bioinformatics analysis

**DOI:** 10.1101/2020.08.18.256735

**Authors:** Hanming Gu, Gongsheng Yuan

## Abstract

There is an urgent need to understand the pathogenesis of the severe acute respiratory syndrome coronavirus clade 2 (SARS-CoV-2) that leads to COVID-19 and respiratory failure. Our study is to discover differentially expressed genes (DEGs) and biological signaling pathways by using a bioinformatics approach to elucidate their potential pathogenesis. The gene expression profiles of the GSE150819 datasets were originally produced using an Illumina NextSeq 500 (Homo sapiens). KEGG (Kyoto Encyclopedia of Genes and Genomes) and GO (Gene Ontology) were utilized to identify functional categories and significant pathways. KEGG and GO results suggested that the Cytokine-cytokine receptor interaction, P53 signaling pathway, and Apoptosis are the main signaling pathways in SARS-CoV-2 infected human bronchial organoids (hBOs). Furthermore, NFKBIA, C3, and CCL20 may be key genes in SARS-CoV-2 infected hBOs. Therefore, our study provides further insights into the therapy of COVID-19.

## Introduction

The coronavirus disease-19 (COVID-19) pandemic caused by severe acute respiratory syndrome coronavirus 2 (SARS-CoV-2) has spread globally (1). Coronaviruses are single-stranded ribonucleic acids (ssRNA) which contain genome sizes ranging from 26 to 32kb (2). SARS-CoV-2 is formed with structural proteins such as spike glycoproteins, membrane glycoproteins, envelope and nucleocapsid proteins, and other non-structural proteins including papain-like proteases and coronavirus main proteases (3). The infection of cells may trigger host immunity to eliminate viral infections (4). However, an insufficient immune response may cause severe damage to patients. Most of the serious infected patients have invasive lesions and are accompanied by inflammation, respiratory distress syndrome, and death (5). This virus may lead to a difficult-to-treat disease which affects not only cilia activities but also wide signaling pathways. Moreover, the human bronchial epithelial cells with ACE2 receptors are the gates for the SARS-CoV-2 invasion (6). Microarray technologies on COVID-19 have been performed in several studies (7, 8). However, the shortcomings of microarray, such as the limited sample size (9), lead to a major challenge to the understanding of key pathogenesis of COVID-19.

In this study, we investigated the effect of SARS-CoV-2 on human bronchial organoids (hBOs) which were generated from primary human bronchial epithelial cells (hBEpC). We analyzed the GEO data (GSE150819), provided by Okamoto T and Takayama K (Osaka University), from the Gene Expression Omnibus database (http://www.ncbi.nlm.nih.gov/geo/) to identify DEGs and the relevant biological processes by utilizing comprehensive bioinformatics analysis. These pathogenetic genes and pathways could be the targets for future therapeutic interventions.

## Methods

### Data resources

Gene expression profile dataset GSE150819 was downloaded from the GEO database (http://www.ncbi.nlm.nih.gov/geo/). The data was produced by using an Illumina NextSeq 500 (Homo sapiens) (Research Institute for microbial diseases, Osaka University, Suita, Osaka, Japan).

### Data acquisition and preprocessing

The raw microarray data files between SARS-CoV-2 infected human bronchial organoids (hBOs) from commercially available cryopreserved primary human bronchial epithelial cells (hBEpC) and uninfected controls were subsequently conducted by R script. We utilized a classical t-test to identify DEGs with P<.05 and fold change ⩾1.5 as being statistically significant.

### Gene Ontology (GO) and pathway enrichment analysis of DEGs

Gene Ontology (GO) analysis and the Kyoto Encyclopedia of Genes and Genomes (KEGG) database are commonly used for systematic analysis of gene functions and annotation of biological pathways. GO analysis and KEGG pathway enrichment analysis of DEGs in this study were performed by the Database for Annotation, Visualization, and Integrated Discovery (DAVID) (http://david.ncifcrf.gov/) online tools. P<0.05 and gene counts >10 were considered statistically significant.

### Module analysis

Molecular Complex Detection (MCODE) was used to detect densely connected regions in Protein-protein interaction (PPI) networks. The significant modules were from the constructed PPI network using MCODE from Cytoscape software (https://cytoscape.org/). the functional and pathway enrichment analyses were performed using DAVID (http://david.ncifcrf.gov/) online tools.

## Results

### Identification of DEGs in SARS-CoV-2 infected hBOs

To understand the mechanism on SARS-CoV-2 infected human bronchus, the modular transcriptional signature of SARS-CoV-2 infected human bronchial organoids (hBOs) was compared to that of the uninfected controls. A total of 89 genes were identified to be differentially expressed in SARS-CoV-2 infected hBOs with the threshold of P<0.01. Among these DEGs, 59 were up-regulated and 30 down-regulated in SARS-CoV-2 infected hBOs compared with the negative controls. The top 10 up- and down-regulated genes for SARS-CoV-2 infected hBOs and negative controls are listed in Table 1.

**Table 1.**

| Entrez gene | Gene Symble | Fold-change | Regulation |
| --- | --- | --- | --- |
| **Top 10 down-regulated DEGs** |  |  |  |
| 282996 | RBM20 | -1.818760446 | Down |
| 85364 | ZCCHC3 | -1.817160181 | Down |
| 10626 | TRIM16 | 1.816969595 | Down |
| 114785 | MBD6 | -1.815479751 | Down |
| 399948 | COLCA1 | -1.814930811 | Down |
| 6330 | SCN4B | -1.814353104 | Down |
| 7710 | ZNF154 | -1.813107418 | Down |
| 23368 | PPP1R13B | -1.812761468 | Down |
| 199920 | C1orf168 | -1.812479327 | Down |
| 1031 | CDKN2C | -1.810848576 | Down |
| **Top 10 up-regulated DEGs** |  |  |  |
| 127544 | RNF19B | 1.823564136 | up |
| 146227 | BEAN1 | 1.822745659 | up |
| 124872 | B4GALNT2 | 1.819974513 | up |
| 2621 | GAS6 | 1.81993098 | up |
| 4502 | MT2A | 1.819527811 | up |
| 3460 | IFNGR2 | 1.819426461 | up |
| 4792 | NFKBIA | 1.81675265 | up |
| 2947 | GSTM3 | 1.816714761 | up |
| 6364 | CCL20 | 1.816339851 | up |
| 10125 | RASGRP1 | 1.815833075 | up |

### Enrichment analysis of DEGs in SARS-CoV-2 infected hBOs

To further analyze the biological roles of the DEGs from negative controls versus SARS-CoV-2 infected hBOs, we performed the KEGG pathway and GO categories enrichment analysis (Figure 1). Gene Ontology (GO) enrichment is used for interpreting sets of genes, in which genes are assigned to a set of predefined areas depending on their functional characteristics (10). We herein identified the top 3 cellular components: “Cell fraction”, “Insoluble fraction”, and “Membrane fraction”. We then identified the top 3 biological processes including “Death”, “Cell death”, and “Apoptosis”. We also identified the top 3 molecular functions: “Enzyme inhibitor activity”, “Peptidase inhibitor activity”, and “Endopeptidase inhibitor activity” (Figure 1).

**Figure 1.**
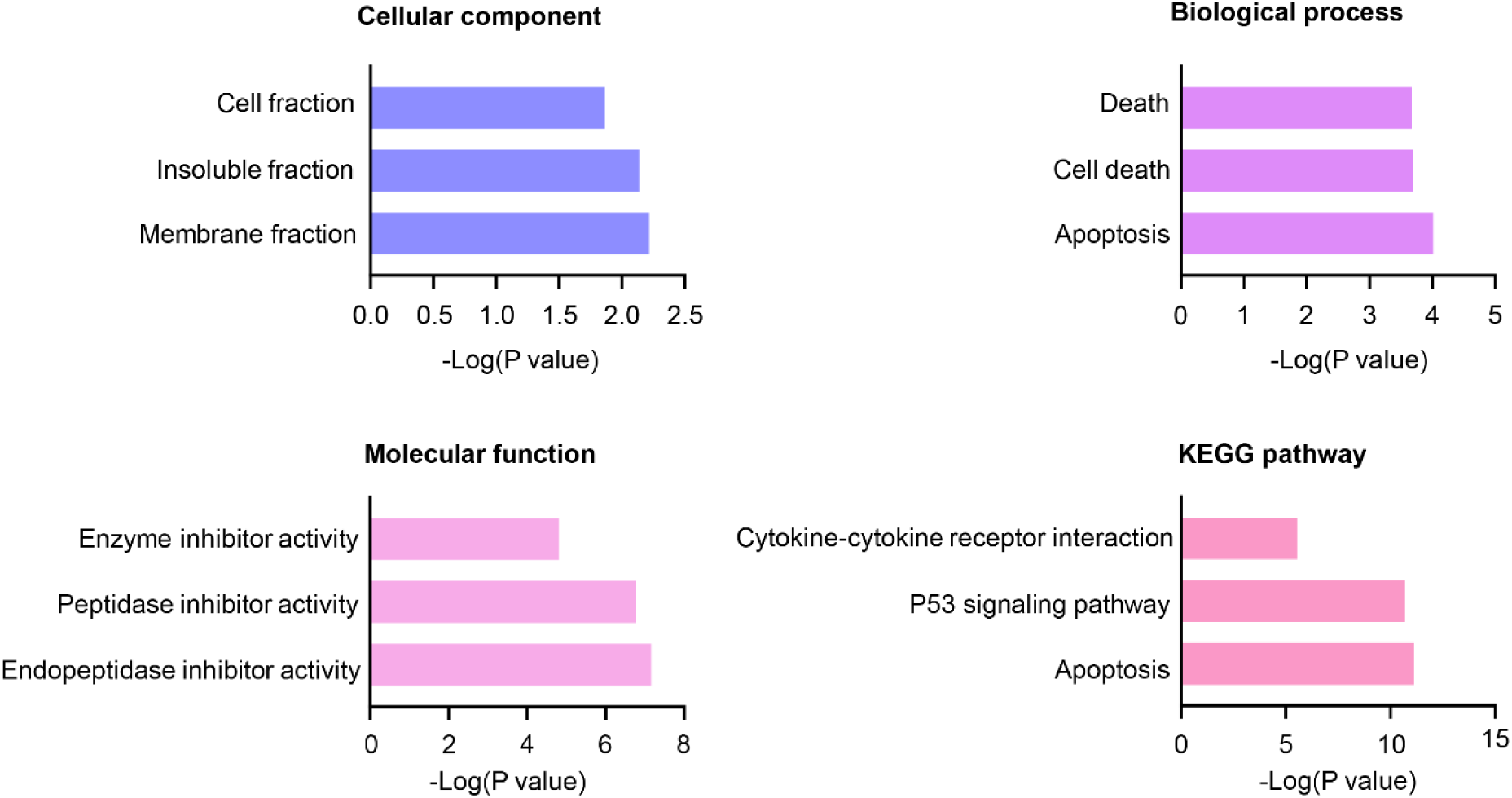
The Cellular component, Biological process, Molecular function terms, and KEGG pathways enriched by the DEGs between SARS-CoV-2 infected hBOs and controls. KEGG = Kyoto Encyclopedia of Genes and Genomes.

KEGG is an integrated database resource and a computationally generated database in many categories including metabolism, other cellular processes, organismal functions, and human diseases (11). In this study, we identified the top 3 KEGG pathways: “Cytokine-cytokine receptor interaction”, “P53 signaling pathway”, and “Apoptosis” (Figure 1).

### PPI network analysis of DEGs in SARS-CoV-2 infected hBOs

To explore the relationships of DEGs, we used the Cytoscape software to set up a PPI network at protein levels. The predefined criterion of combined score >0.7, a total of 30 interactions, and 84 nodes were created to form a PPI network between SARS-CoV-2 infected hBOs and the uninfected hBOs. The top 5 hub genes with degree scores are depicted in Table 2.

**Table 2.** Top five genes demonstrated by connectivity degree in the PPI network

| Gene Symbol | Gene title | Degree |
| --- | --- | --- |
| NFKBIA | NFKB inhibitor alpha | 6 |
| C3 | complement C3 | 5 |
| CCL20 | C-C motif chemokine ligand 20 | 4 |
| BCL2A1 | BCL2 related protein A1 | 3 |
| BID | BH3 interacting domain death agonist | 3 |

### Module analysis

The top significant modules of SARS-CoV-2 infected hBOs versus uninfected controls were analyzed by the functional annotation of the genes (Figure 2). The functional genes were then analyzed by String and DAVID. We identified the top ten significant biological processes or pathways: “Leishmaniasis”, “Apoptosis”, “Measles”, “Tuberculosis”, “Kaposi’s sarcoma-associated herpesvirus infection”, “Legionellosis”, “P53 signaling pathway”, “Pertussis”, “NF-kappa B signaling pathway”, and “Chagas disease (American trypanosomiasis)” in Supplemental Table S1.

**Figure 2.**
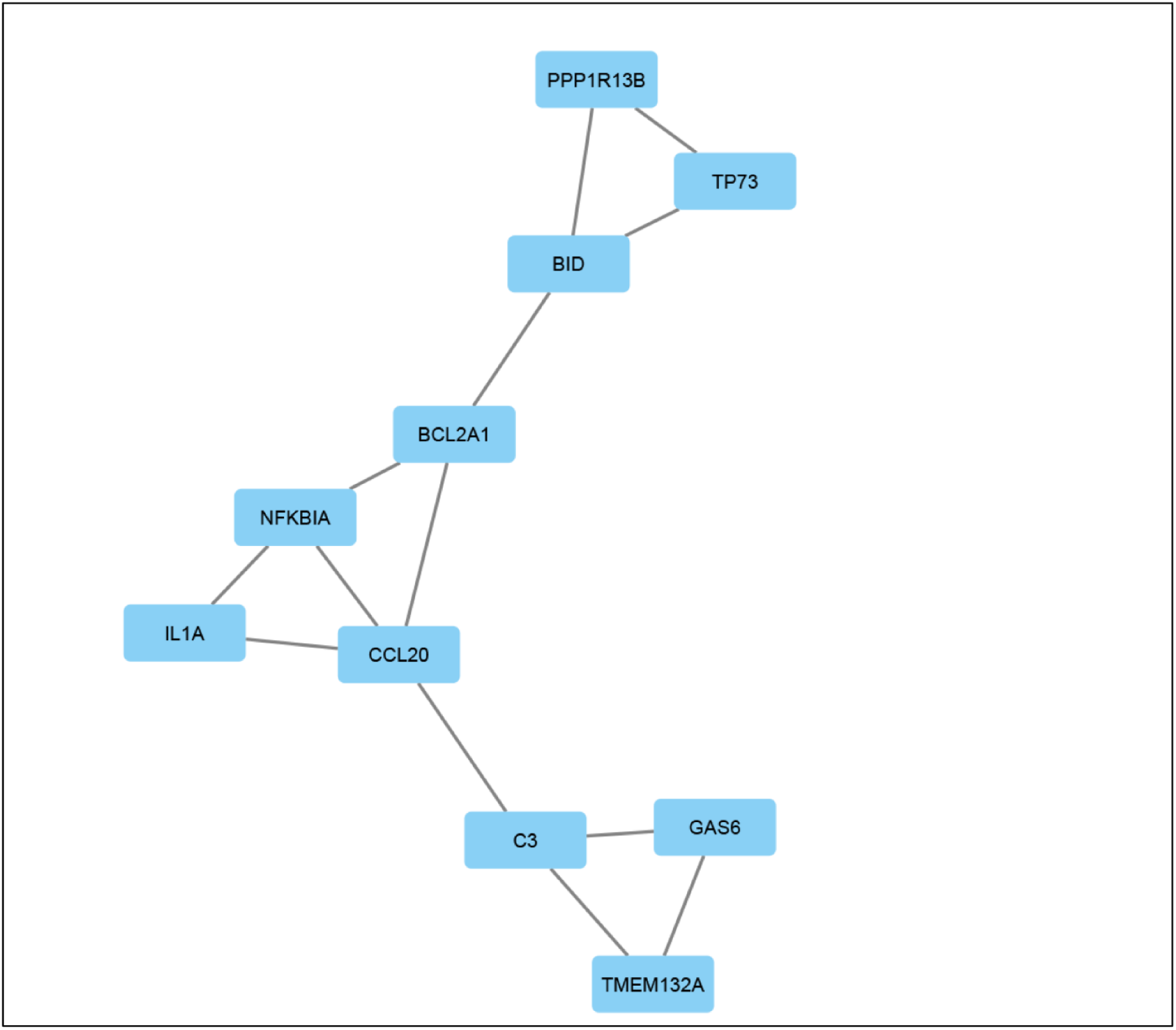
Top module from the protein-protein interaction network between SARS-CoV-2 infected hBOs and controls.

## Discussion

COVID-19 is a highly infectious and transmissible disease among humans (12). Small groups of patients with COVID-19 who experience severe pneumonia can cause ARDS and death (13). The primary step after inhalation of SARS coronaviruses is an invasion of epithelial cells through binding the SARS spike protein to angiotensin-converting enzyme 2 (ACE2) receptors (14). An important mechanism regulating lung pathology in SARS infections is a cytokine storm leading to the “inflammation storm” or “macrophage activation syndrome” (15). However, how SARS-CoV-2 affected the signaling pathways and potential mechanisms in epithelial cells or bronchus is totally unknown.

In our study, we identified 59 up-regulated and 30 down-regulated DEGs between SARS-CoV-2 infected hBOs and the control ones. To better understand the DEGs, we first performed the GO analysis. Interestingly, we found the death and apoptosis process were mainly shown in the biological process panel. A recent report showed that XAF1, TNNF, and FAS induce T cell apoptosis in COVID-19 patients (16). The previous report showed cilium abnormalities in bronchus are associated with COVID-19 (17). The cilium shrinking and abnormal beating lead to secondary infections, apoptosis, and cell death (18). Also, apoptosis and pericyte loss are identified in alveolar capillaries in COVID-19 infections (19). Cell death in response to oxidative stress is also found in the global analysis of SARS-CoV-2 protein interactions (20). Similarly, the apoptosis and death signaling are critical steps when SARS-CoV-2 infected hBOs. It is believed that, during the progress of COVID-19, SARS-CoV-2 will infect extensively tracheal cells leading to apoptosis and necrosis. COVID-19 is involved in the regulation of enzyme activities in hBOs (21). During the synthesis of SARS-CoV-2, the polyproteins are translated from the viral RNA (22). We found that “Enzyme inhibitor activity”, “Peptidase inhibitor activity”, and “Endopeptidase inhibitor activity” were the top three processes in SARS-CoV-2 infected hBOs. These findings indicated that SARS-CoV-2 may inhibit the enzyme inhibitor activity in hBOs to create more polyproteins and viruses. Thus, hBOs may be a potential virus production base during COVID-19 diseases.

KEGG analysis showed the “Cytokine-cytokine receptor interaction” is the main pathway in SARS-CoV-2 infected hBOs. Cytokines such as IL-1, IL-6, and TNFα are in response to an activating stimulus, and they trigger the cell response via association with the specific receptors on the cell surface (23). Interestingly, the SARS-CoV-2 spike (S) protein binds angiotensin-converting enzyme 2 (ACE2) and promotes cellular entry (24). Thus, our results suggested that SARS-CoV-2 entered cells accompanying with many cytokines binding reactions. To identify these cytokines and inhibit the bindings may become an effective strategy to treat COVID-19 diseases.

After analyzing the PPI network, we found the NF-κB pathway is involved in the COVID-19. The NF-κB pathway is considered as an inflammation center, which is based on the role in the expression of proinflammatory genes including cytokines, chemokines, and adhesion molecules (25). NF-κB activation induces various target genes in inflammatory diseases and NF-κB signaling crosstalk affects many signaling pathways such as STAT3, RGS12, WNT–β-catenin, and P53 (26-28). Cytokine storm syndromes and immunosuppression, which are mainly regulated by NF-κB, are the specific pathological processes in the severe COVID-19 patients (29, 30). Thus, inhibition of NF-κB therapy can inhibit the infection caused by SARS-CoV-2.

In summary, NFKBIA, C3, and CCL20 may be key genes in SARS-CoV-2 infected hBOs. Our study suggested that “Cytokine-cytokine receptor interaction”, “P53 signaling pathway”, and “Apoptosis” were the main signaling pathways during SARS-CoV-2 infection. Our data provide further evidence for the intervention during SARS-CoV-2 infection.

